# Belowground competition favors character convergence but not character displacement in root traits

**DOI:** 10.1101/2020.08.14.251280

**Authors:** Sara M Colom, Regina S Baucom

## Abstract

- Character displacement can play a major role in species ecology and evolution, however, research testing whether character displacement can influence the evolution of root traits in plant systems remains scarce in the literature. Here we investigated the potential that character displacement may influence the evolution of root traits using two closely related morning glory species, *Ipomoea purpurea* and *I. hederacea*.
- We performed a field experiment where we grew the common morning glory, *I. purpurea*, in the presence and absence of competition from *I. hederacea* and examined the potential that the process of character displacement could influence the evolution of root traits.
- We found maternal line variation in root phenotypes and evidence that belowground competition acts as an agent of selection on these traits. Our test of character displacement, however, showed evidence of character *convergenc*e on our measure of root architecture rather than *displacement*. These results suggest that plants may be constrained by their local environments to express a phenotype that enhances fitness. Therefore, the conditions of the competitive environment experienced by a plant may influence the potential for character convergence or displacement to influence the evolution of root traits.

## Introduction

Character displacement has long been considered an important mechanism that may facilitate species coexistence, result in the evolution of novel phenotypes, and potentially promote the adaptive radiation of species (Brown & Wilson, 1956; Losos 2000; Pfennig *et al*., 2006; Pfennig & Pfennig 2009). Character displacement is hypothesized to occur due to high phenotypic similarity between related species, which leads to increased competition and a concomitant reduction in fitness (Schluter 2000; Pritchard & Schluter 2001; Day & Young 2004; Dayan & Simberloff 2005). Despite the importance of character displacement in evolutionary processes, the majority of character displacement work has been performed in animal systems (Schluter & Grant 1984; Schluter *et al*.,1985; Losos 1990 & 2009; Schluter & McPhail 1992; Pritchard 1998; Martin & Pfennig 2011), such that comparatively fewer well-supported studies of character displacement exist in plants (Levin 1985; Muchhala & Potts 2007; Muchhala 2008; Hopkins & Rausher 2011; Beans, 2014).

Of the available research on character displacement in plants, the majority is focused on the evolution of floral morphology and color as a response to competition for pollinators (Armbruster 1985 & 1986; Muchhala & Potts 2007; Muchhala 2008; Smith & Rausher 2008; Beans 2014). Other plant traits contribute to fitness and mediate plant-plant competition, however, and these remain largely unstudied as targets of character displacement. For example, the belowground root system plays a critical role in the acquisition of minerals and water from the soil (Fitter,1987; Fitter, 2002) and in mediating belowground plant-plant competition (Casper and Jackson 1997, Kroon *et al*., 2003 & Schenk 2006; Ravenek *et al*., 2016). Thus, the belowground root system could likely respond to selection *via* competition between closely related, co-occurring species, potentially leading to character displacement in root traits. However, research examining the potential for character displacement on the root system remains a significant gap in evolutionary ecology.

The belowground root system is a complex organ composed of diverse and developmentally interdependent traits that are often cataloged into ‘functional groups’ that influence resource uptake in different ways (Fig. 1; Fitter 1987; Lynch 1995; Bucksch *et al*., 2014). Phenotypic diversity in the traits within functional groups can influence how plants explore the soil and acquire nutrients – *e*.*g*., the morphology of individual root traits (*e*.*g*., root diameter), how components of traits are arranged spatially (architecture, *e*.*g*., angles), the size of these traits and the volume they take up (root system size) and the distribution of these elements over space (root system topology; Fig. 1; Lynch & Brown, 2001; Fitter *et al*., 2001; Lynch 2005; Nguyen & Stangoulis 2019; Canales *et al*., 2019). Despite the important ecological and functional role of the root system, only a few studies have explicitly investigated the potential that natural selection can lead to the phenotypic evolution of root traits (Ferguson *et al*., 2016; Murren *et al*., 2020; Colom & Baucom 2020), and even fewer have considered the role that belowground competition may play in the evolution of the root system (Colom &Baucom 2020).

**Fig. 1.**
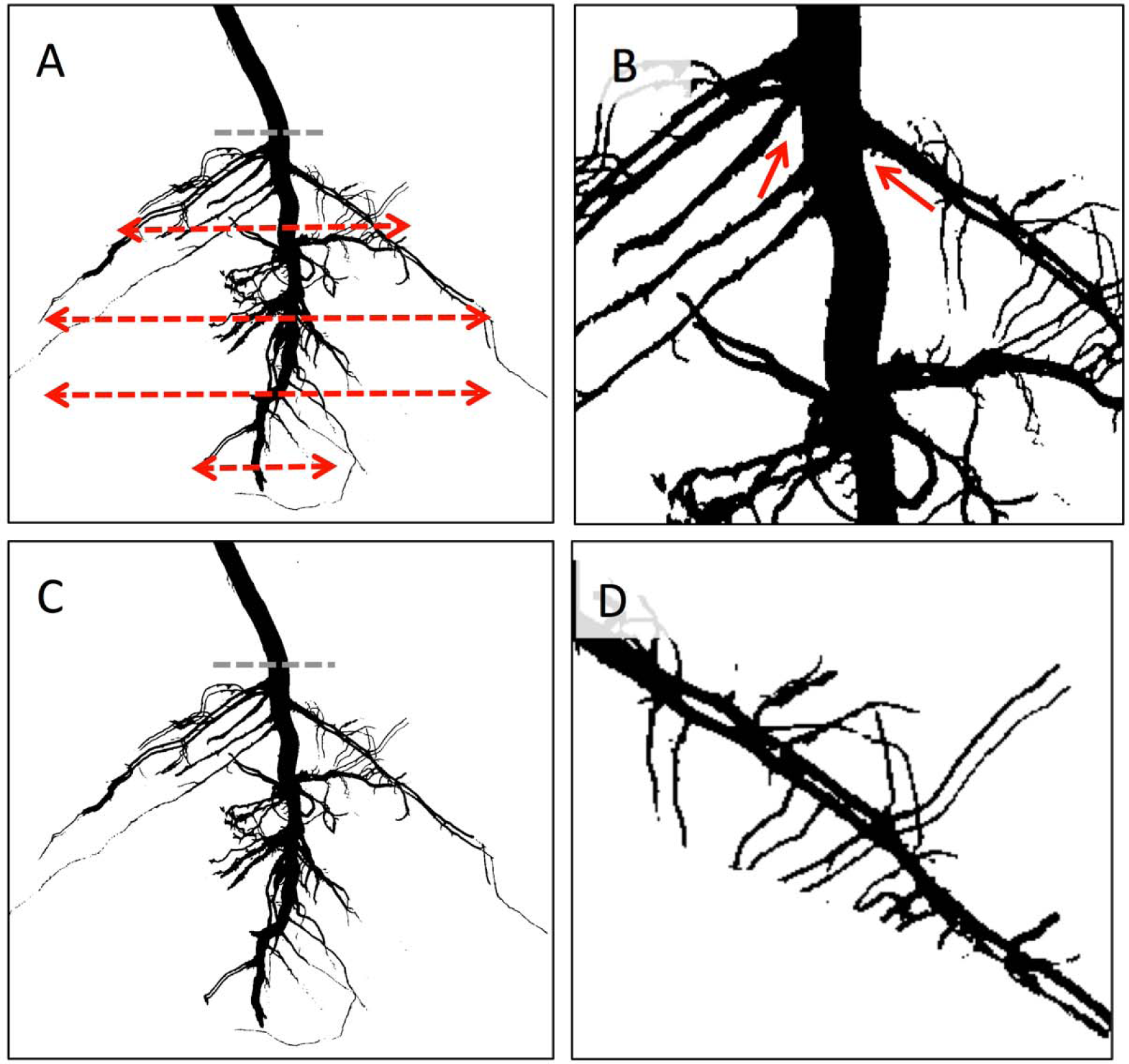
A depiction of four distinct functional root trait classes, (A) root topology, (B) architecture, (C) size, and (D) morphology. *Root topology* describes the general shape of the root system--*e*.*g*., root system width with soil depth as indicated with the dashed red arrows of varying lengths beneath the soil line, shown by the grey dashed line. *Root architecture* is a suite of traits that describe the spatial arrangement of the root system including root angle formed between tissues (‘Root tissue angle’, indicated with the red arrows), overall root system width and length, and branching patterns (distances between lateral root nodes). *Root size* encompasses root traits such as root surface area and volume of the root system beneath the soil line (indicated with grey dashed line). *Root morphology* is a suite of traits that describes characteristics of individual root traits (*e*.*g*., lateral root length, root diameter), and the relative number of individual root traits such as lateral root number and diameter. These traits are emphasized here with a close up depiction of a single lateral root where root length, diameter and number of second order branching roots are more readily visible.

Previously, we demonstrated that competitive belowground interactions can act as an agent of selection on root traits in *Ipomoea purpurea* and *I. hederacea*, two closely related species of morning glory that are found to co-occur, and compete, in agricultural fields and other areas of high disturbance (Colom &Baucom 2020). We found evidence of genetic variation underlying traits associated with both root morphology and architecture (*i*.*e*., primary root length, angle and width), and additionally showed that competition between the two species influenced the pattern of selection on root traits. Specifically, when in competition with *I. purpurea, I. hederacea* individuals with shallower root architectures – a trait associated with increased topsoil foraging (Fitter 1987; Lynch 1995) – exhibited higher fitness. In contrast, belowground competition from *I. hederacea* altered the pattern of selection on root system size in *I. purpurea*. When *I. purpurea* was grown in the absence of *I. hederacea*, selection favored individuals with larger root systems, whereas there was no detectable selection on root system size in *I. purpurea* in the presence of competition with *I. hederacea* (Colom &Baucom, 2020). These findings indicate that root traits can respond to selection, and that belowground competition acts as an agent of selection, potentially influencing the evolution of root traits.

Here, we ask “Can belowground competition between closely related species potentially result in character displacement of their root system traits?” We addressed this question by growing maternal lines of *I. purpurea* in the presence and absence of *I. hederacea* and determining if there is a relationship between fitness and the phenotypic distance of multivariate measures of root topology, architecture, size and morphology between these two species. We adopted part of the criteria used to evaluate evidence for the pattern of character displacement (McPhail &Schluter, 1992; Losos 2000) and focused on testing evidence for the process of character displacement. We examined the following core components of character displacement: Criterion 1) belowground competition influences fitness, Criterion 2) traits under selection must have a genetic basis, Criterion 3) belowground competition generates non-random fitness differences as a function of phenotypic variation (*i*.*e*., competition is the agent of selection on phenotype), and Criterion 4) when in competition, the fitness of individuals increases with greater phenotypic distance in root traits compared to their competitor (as depicted in Fig. 2). Criterion 4 is the hallmark prediction of character displacement.

**Fig. 2.**
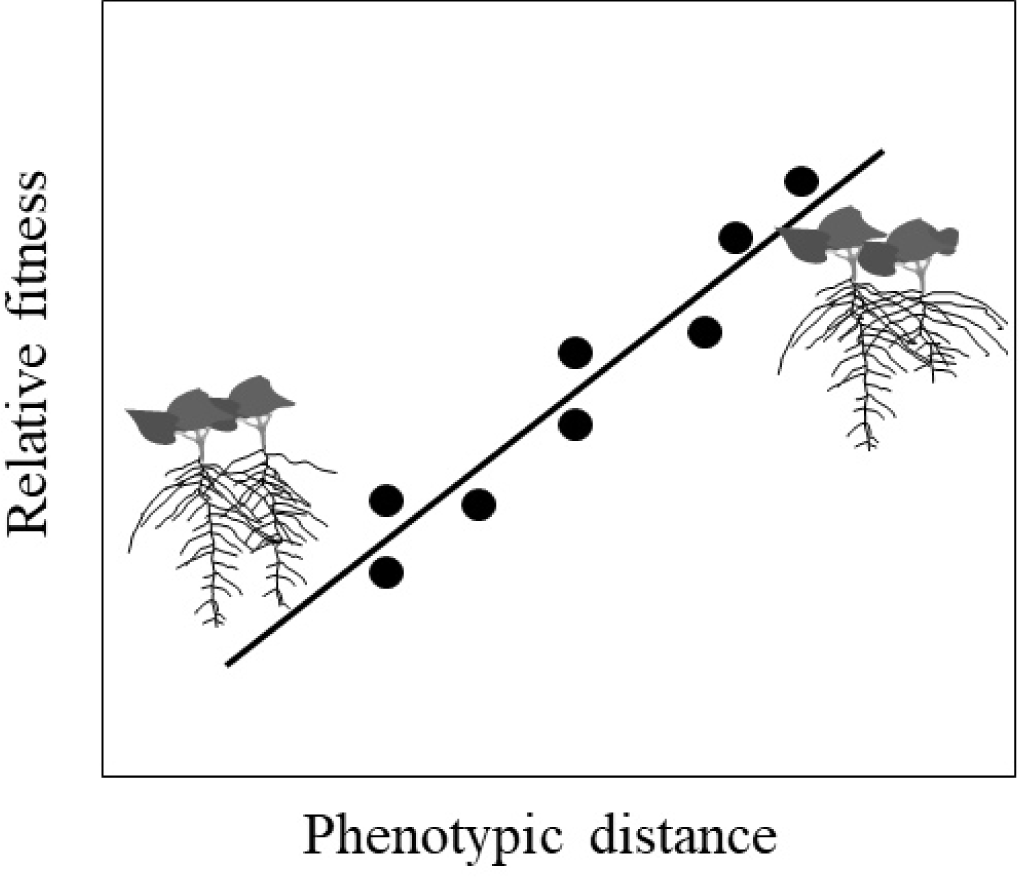
A positive linear relationship between phenotypic distance of competitors’ root traits and relative fitness would support the hypothesis that character displacement can influence the evolution of root traits. Superimposed on the plot are sketches of two pairs of competitors with low (left pair) and high (right pair) phenotypic distance, respectively.

## Materials and Methods

### Study system

We used the closely related morning glory species, *I. purpurea* (L.) Roth and *I. hederacea* (L.) Jacquin (Convolvulaceae) as our experimental system. These *Ipomoea* species commonly co-occur in eastern United States (RS Baucom, *pers. obs*.) and historical records indicate their presence as early as 1700’s and mid 1800 for *I. purpurea* and *I. hederacea*, respectively (Pursh, 1814; Bright, 1998). These *Ipomoea* species are annual, self compatible weedy climbing vines that reside in similar habitats and exhibit high within species morphological diversity in aboveground and belowground traits (Baucom *et al*., 2011; Colom & Baucom, *pers. obs*.). Both species germinate between the months of May and August, begin to flower about six weeks after germination and continue to flower until they are killed at first frost. In this experiment, we used *I. purpurea* as our focal species to have a sufficiently high number o replicates to examine the potential for maternal line and while maintaining a feasibly sized experiment.

### Field design and planting

We obtained ten maternal lines of *I. purpurea*, and six maternal lines of *I. hederacea* from a single population located in Pennsylvania and selfed them for one generation in greenhouse conditions to reduce maternal effects. We planted once-selfed seeds of *I. purpurea* in both the presence of interspecific competition with *I. hederacea*, hereafter, ‘competition’ treatment, and in the absence of competition, hereafter, ‘alone’ treatment at a 42m × 40m field plot located at the Matthaei Botanical Gardens, Ann Arbor, MI on June 2nd of 2018. The field plot was tilled one week prior to planting. For the competition treatment, we used ten maternal lines of *I. purpurea* paired with each combination of six maternal lines of *I. hederacea* to yield 60 ‘unique combination pairings’. We replicated each of these pairings sixteen times. For the alone treatment, we replicated each *I. purpurea* maternal line sixteen times.

All seeds were planted in a complete random block design in which we arrayed four replicates of each unique pairing and maternal lines grown alone in each of four 10 m × 30 m spatial blocks. Blocks were separated from each other by two meters, and seeds planted in the competition treatment were placed three inches away from each other. All plant pairs in the competition treatment and individual plants in the alone treatments were planted 1 m^2^ apart from one another. We placed 1 m tall bamboo stakes next to each experimental seedling and later trained them to grow onto the bamboo sticks to prevent experimental plants from tangling and competing aboveground, following Colom and Baucom, 2020. Because our focal species in this experiment was *I. purpurea*, we centered our data collection and analysis on this species with *I. hederacea* as the closely related competitor.

We watered seeds and recorded germination daily for the first two weeks, and subsequently, relied on rainfall to water our plants. Vole herbivory, which killed some plants during the course of the experiment, was recorded. We kept the soil within a six-inch radius around each experimental plant free of weeds and removed any non-experimental morning glories from the field. One month after planting we counted leaves of each *I. purpurea* plant to serve as a proxy of plant size.

### Root excavation and phenotyping

When experimental plants were reproductively mature, we excavated a subsample of individuals to obtain root phenotype data from individuals grown in each treatment. We sampled between four to eight biological replicates of each maternal line of *I. purpurea* in the alone treatment, and four to eight biological replicates of *I. purpurea* and *I. hederacea* planted in competition for each of the unique 60 combination pairings. In total we excavated and phenotyped 511 plants. We adopted the shovelomics method for root excavation (Colombi *et al*., 2015) as previously described (Colom & Baucom, 2020) and imaged their roots with a high-resolution camera, Canon EOS Rebel XSi 12.2 MP (18-55mm IS Lens).

Each of the images was imported to DIRT (Das *et al*., 2015), a fully automated online program designed to phenotype multiple root traits from plants sampled in field conditions. We removed traits from this output that were not applicable for our study system, such as monocot root traits, as well as highly redundant traits (*i*.*e*., represented a mathematical combination of two or more traits). All trait measurements computed by DIRT rely on estimates of root length, diameters, branching angles, density and spatial root distribution that are quantified from the pixels of an image mask of the root system (binarized image of the root system) and a structural description of the root system or ‘skeleton’. The structural description (‘root tip path’ in DIRT) of the root system is a curve representation of the root system based on different samples points that allows the program to capture multiple measurements (‘skeleton’ traits in DIRT) that are otherwise occluded or confounded—*e*.*g*., the network of a mature root system occludes its interior and smaller roots may bind together and appear as a single root.

### Specific root traits analyzed

We examined a total of 33 traits, which we *a priori* classified into the four functional classes of root architecture, morphology, size and topology, *e*.*g*., root angle and horizontal/vertical length, lateral root number and diameter, total root system surface area and maximum width of the root system for a given soil depth, respectively (*see* Table S2-1 & Fig. S2-1).

### Fitness data

We began to collect mature fruit of experimental *I. purpurea* in September and continued to do so until all plants senesced in mid-October. We sampled between three to eight replicates for each maternal line per *I. purpurea* in the alone treatment, and between two and nine replicates of *I. purpurea* for each competition pairing; in total we sampled seed from 429 *I. purpurea* (Num. alone = 62 & Num. in competition = 367).

## Statistical analysis

All statistical analyses were performed in R version 3.0 (R core team 2018).

### Modular root traits

We elected to perform a principal component analysis (PCA) to reduce the high dimensionality of our root phenotypes (see *Root system traits* section) with the correlation matrix of 33 root traits rendered by DIRT (Table S1) using the ‘PCA’ function from the ‘factoMiner’ package (Le *et al*., 2008). Prior to PCA we mean centered each trait to zero, scaled the standard deviation to a value of one, and applied a box cox transformation to reduce skewness in the data with the ‘preProcess’ function of the ‘caret’ package (Khun, 2019). Since preliminary visualization of the PC’s showed grouping by block, we performed the PCA on the indexed residuals of each root trait after controlling for block in a one-way ANOVA. A scree plot performed on the output of this PCA showed that each of the first four PC’s explained at least 10.0% of the total phenotypic variation (Fig. 3). Therefore, we focused our analysis on the first four PC’s as modular root traits. To evaluate how each individual root trait contributed to PC1-4, we calculated the proportion of squared loading coefficients to the sum of squares with the ‘fviz_contrib’ function in the ‘factoextra’ package (*see* Fig 2; Kassambara & Mundt, 2017. We found that the first four PCs were associated generally by traits that describe topological, architectural, size and morphological aspects of the root system, respectively, (*see* Fig. 3; Table S1). We hereafter refer to PC1-4 as root topology, architecture, size and morphology, respectively.

**Fig. 3.**
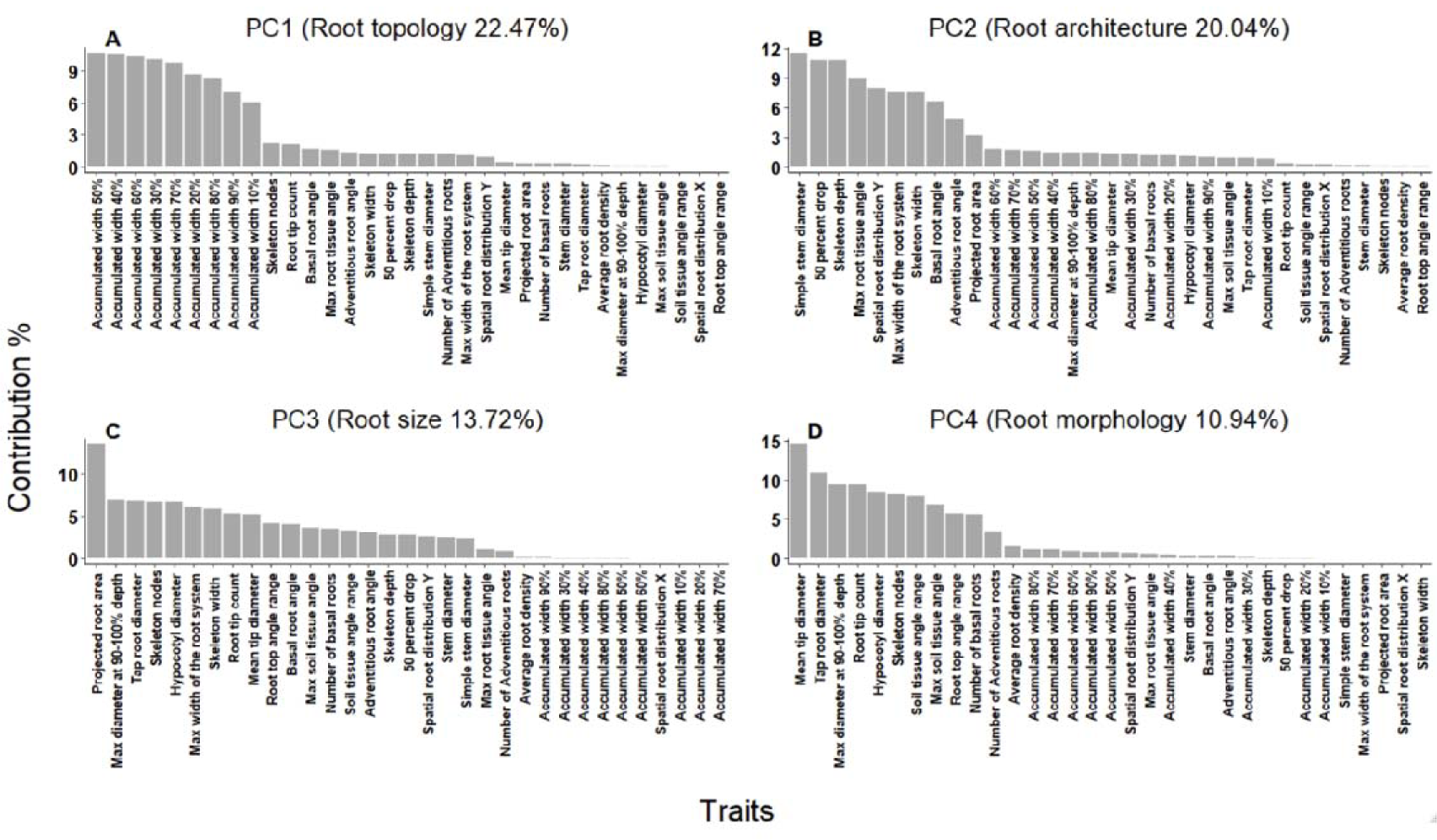
Bar graphs demonstrating the contribution of individual root traits to the first four PC’s (A-D) after removing Block effects. We refer to these four PC’s as topology (PC1), root architecture (PC2), root size (PC3) and root morphology (PC4). Individual traits that contribute to each modular root trait are defined in Table S1.

### Evidence of belowground competition

To test whether belowground competition influences fitness, we performed a linear mixed model where we used the observed seed number as our response variable, block, treatment and block by treatment as fixed effects, and maternal line and treatment by maternal line interaction as a random effects. We excluded treatment by maternal line interaction in our final model because we found that its inclusion did not improve akaike information criterion (AIC) when we compared it to a model that lacked this interaction term. Because preliminary analysis showed a strong correlation between leaf number, a proxy for plant size, and seed number, we also included leaf number as a covariate in our model. We did *F*-tests with type three sums of squares using Satterthwaite’s method to evaluate the significance of fixed effects, and log-likelihood ratio χ2 tests to test for the random effect using the ‘anova’ and ‘ranova’ functions of the ‘lmerTest’ package (Kuznetsova *et al*., 2017). We estimated the least square means of seed number for each treatment averaged across block and block by treatment interaction with the ‘emmeans’ function as above (Length, 2019).

### Maternal line variation of root traits

To determine if there was evidence of maternal line variation in modular root traits of plants grown in the field, we performed separate linear mixed models for each PC’s. We ran separate models for each PC as block, treatment and treatment by block interaction as fixed effects, and maternal line and maternal line by treatment interaction were random effects. Because we found that maternal line by treatment and block by treatment interactions did not improve the AIC when we compared it to a model that lacked these interaction terms, we removed these factors from our final models. We performed *F*-tests and log-likelihood ratio χ2 tests as above to assess the significance of fixed and random effects, respectively. We also evaluated evidence for block, treatment and maternal line variation on eight individual root morphology traits and architecture traits *post hoc* within *I. purpurea* because we detected evidence for selection on these traits (see *Selection on root traits* below).

### Calculating standardized relative fitness

To test for selection on root traits and that fitness increases with phenotypic distance, we used standardized relative fitness as our response variable. For our calculation of standardized relative fitness, we divided the observed seed number by the mean seed number for *I. purpurea*, within each competition treatment, (*e*.*g*.,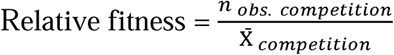, where *n* represents the number of observed seeds from each individual in competition, and X□ represents the mean seed number of plants in competition). Then we averaged the output by maternal line and treatment for the selection analysis and averaged the output by maternal line and combination pairing for the test that fitness increases with phenotypic distance. Then we standardized values of average relative fitness to control for block and plant size by using the residuals of a two-way ANOVA that included only block and leaf number as explanatory variables of average relative fitness.

### Testing for selection on root traits

We performed genotypic selection analysis (Rausher, 1992) to examine if competition imposes selection on modular root traits. We averaged the PC scores (block standardized) of root architecture, topology and morphology by maternal line and treatment, and then performed regressions for each modular trait onto standardized relative fitness (averaged by treatment and maternal line) for each treatment separately. We elected to exclude root size (PC3) from this and subsequent analysis because we did not find evidence for maternal line variation or directional selection on this trait in the presence of competition in previous work (Colom & Baucom 2020), or in the preliminary analysis of the present research. Preliminary assessment of quadratic selection on individual PCs did not reveal evidence for either stabilizing or divergent selection, and thus we report only linear terms. To test whether the pattern of directional selection differed between treatments, we performed ANCOVAs on relative fitness for each PC, wherein treatment, trait and treatment by trait interaction were included as our explanatory variables.

Because not all root traits necessarily contribute to fitness, the effect of selection on any individual trait contributing to a PC can be obscured (Mitchell-Olds & Shaw 1987; Chong *et al*., 2018). Therefore, we performed ‘PC back regression’, which is a linear transformation technique where we can input PC’s of interest and their corresponding eigenvectors to recover the selection gradients acting on specific root traits in their original trait space. Specifically, selection gradients on the original root traits are reconstructed by projecting the regression coefficients from our selection analysis onto their corresponding eigenvectors (Jolliffe 2002, p. 169; Chong *et al*., 2019). We used a matrix with the eigenvectors of *modular* root topology, architecture and morphology, standardized for block, and a vector of their corresponding selection gradients (R script available in supplementary). We performed matrix multiplication as shown by the formula, β = *EA*, where β represents a vector of the reconstructed selection gradients on the original root traits, *E* is the matrix of the three PC’s eigenvectors standardized by block, and *A* is a vector of the regression coefficients obtained from regressing relative fitness on these PC scores. We calculated reconstructed βs for individuals of *I. purpurea* grown alone and in competition separately. To test if these selection gradients were significantly different from zero, we calculated a standard error for each reconstructed trait by taking the square root of the difference between the squared standard errors obtained from the selection analysis on the PC scores for each treatment. Then, we estimated confidence intervals for each β at an alpha of 0.05% based on plus or minus two standard errors from each β. If the confidence interval of β did not include zero, we interpreted those slopes as different from zero (Table 2). We interpreted selection gradients from one treatment (*i*.*e*. presence or absence of competition) that did not lie within the 95% confidence interval of the other treatment as evidence that belowground competition imposes selection on that trait. We compared slopes in this manner since PC back regression method applied to a subspace results in a loss of information and consequently impacts our ability to estimate the degrees of freedom required for the traditional approach to compare selection gradients, *i*.*e*., ANCOVA F-statistic to calculate its corresponding *F-statistics* (J. Stinchcombe, pers. comm).

**Table 2.**
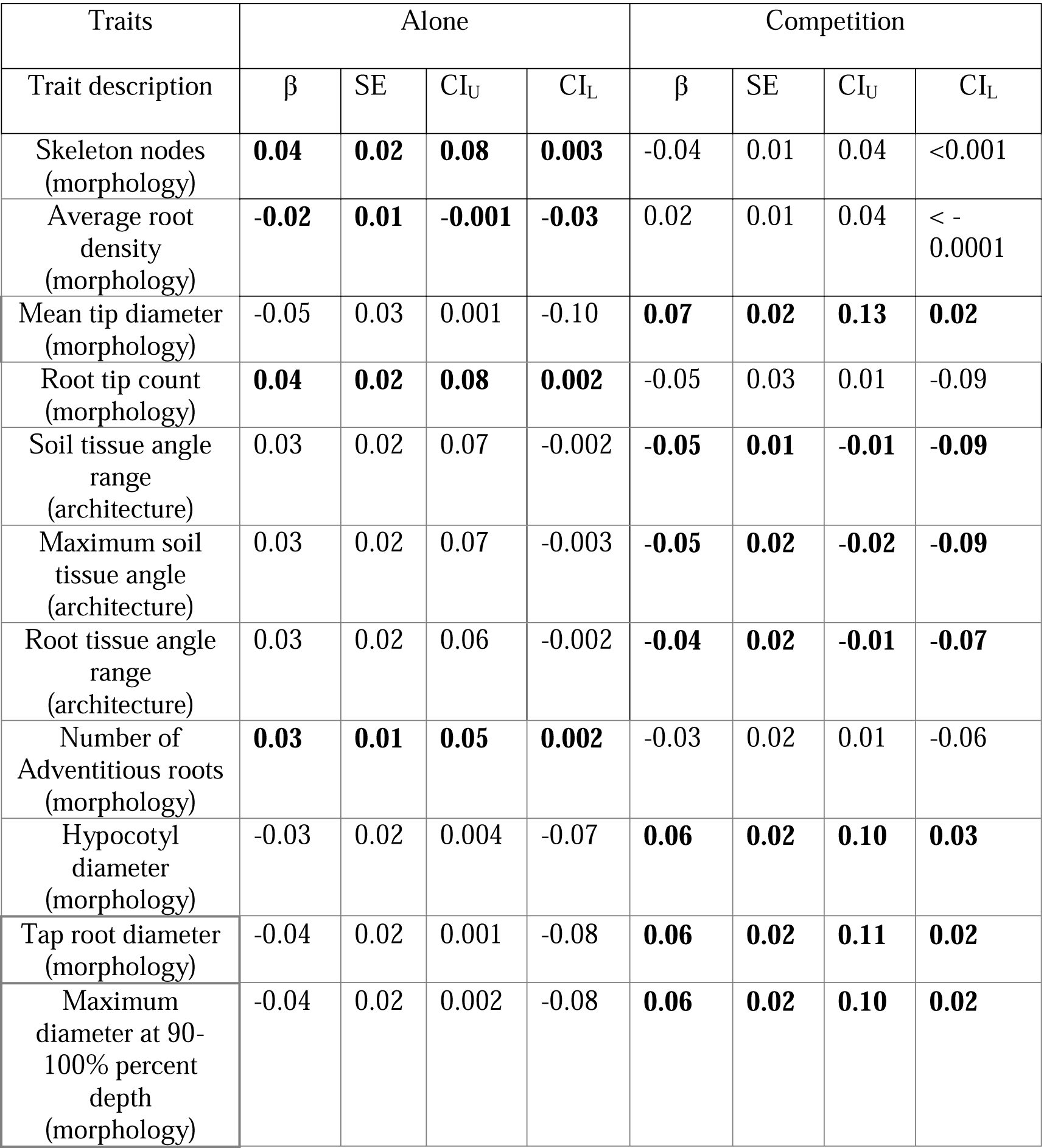
Results of PC back regression of selection gradient projected back onto original traits (β) and corresponding ± 1 standard error (SE) and upper and lower confidence intervals (CI) based on β ± 1.96 × SE. Bolded values indicate a confidence interval of 95% for β that does not include zero. Each root trait was cataloged into four functional trait classes indicated within parenthesis *a priori*.

| Traits | Alone |  |  |  | Competition |  |  |  |
| --- | --- | --- | --- | --- | --- | --- | --- | --- |
| Trait description | $\beta$ | SE | CI<sub>U</sub> | CI<sub>L</sub> | $\beta$ | SE | CI<sub>U</sub> | CI<sub>L</sub> |
| Skeleton nodes (morphology) | **0.04** | **0.02** | **0.08** | **0.003** | -0.04 | 0.01 | 0.04 | <0.001 |
| Average root density (morphology) | **-0.02** | **0.01** | **-0.001** | **-0.03** | 0.02 | 0.01 | 0.04 | < -0.0001 |
| Mean tip diameter (morphology) | -0.05 | 0.03 | 0.001 | -0.10 | **0.07** | **0.02** | **0.13** | **0.02** |
| Root tip count (morphology) | **0.04** | **0.02** | **0.08** | **0.002** | -0.05 | 0.03 | 0.01 | -0.09 |
| Soil tissue angle range (architecture) | 0.03 | 0.02 | 0.07 | -0.002 | **-0.05** | **0.01** | **-0.01** | **-0.09** |
| Maximum soil tissue angle (architecture) | 0.03 | 0.02 | 0.07 | -0.003 | **-0.05** | **0.02** | **-0.02** | **-0.09** |
| Root tissue angle range (architecture) | 0.03 | 0.02 | 0.06 | -0.002 | **-0.04** | **0.02** | **-0.01** | **-0.07** |
| Number of Adventitious roots (morphology) | **0.03** | **0.01** | **0.05** | **0.002** | -0.03 | 0.02 | 0.01 | -0.06 |
| Hypocotyl diameter (morphology) | -0.03 | 0.02 | 0.004 | -0.07 | **0.06** | **0.02** | **0.10** | **0.03** |
| Tap root diameter (morphology) | -0.04 | 0.02 | 0.001 | -0.08 | **0.06** | **0.02** | **0.11** | **0.02** |
| Maximum diameter at 90-100% percent depth (morphology) | -0.04 | 0.02 | 0.002 | -0.08 | **0.06** | **0.02** | **0.10** | **0.02** |

### Fitness increases with phenotypic distance

To test whether the phenotypic distance between root system traits between competitors is positively associated with fitness in *I. purpurea*, we regressed *I. purpurea* standardized relative fitness on the phenotypic distance between root traits of competing plants. For each PC we calculated the Euclidean distances between competitors with the ‘cdist’ function from the ‘rdist’ R package (Blaser, 2018). The calculation of phenotypic distance was done by finding the linear distance within the same modular root system traits: 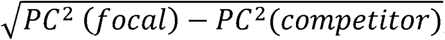, where *PC* represents the prinicpal component of a given axis), for phenotypic distances of root topology, architecture and morphology respectively). We also evaluated phenotypic distance between different types of traits*—e*.*g*., phenotypic distance between PC1 and PC2*—*but did not uncover any evidence that different trait combinations influenced fitness, and thus we do not include these results here. Each metric of phenotypic distance was averaged according to each combination pairing and regressed onto our values of standardized relative fitness (averaged by maternal line, treatment and combination). We used *F*-statistics to ascertain whether the slopes were significantly different from zero.

We also evaluated evidence for the prediction of character displacement on eight individual root morphology traits and three individual root architecture traits *post hoc* as above because we detected evidence for selection on these traits at the individual level, see *Selection on root traits* above.

Because we had multiple biological replicates per sample point in this analysis (N = 2-6 combination pairings), and our main goal was to examine changes in relative fitness given phenotypic distance in root traits, we elected to retain all samples to evaluate the relationship between fitness and phenotypic distance of modular and specific root traits. We also retained two pairings that may be outliers because we found that they had a low amount of variation around the mean (*see* Fig. S4), indicating that these points are not biased by an extreme phenotypic value.

## Results

### Describing the root system as modular root traits

PCA showed that the first four PC’s contributed to 22.5%, 20.0%, 13.7% and 10.9% of the total variation, respectively. Because the traits driving the variation in PC1, PC2, PC3 and PC4 were mainly descriptors of root topology, architecture, system size and morphology, respectively, we refer to them as corresponding modular root traits. For PC1 we found that accumulated root width per soil depth explained most of the variation on this axis (‘root topology’; Fig. 3A). Since each measure of accumulated root width per soil depth loaded positively on this axis, higher scores correspond to a root system with greater root width per soil depth.

Higher PC2 scores were associated with broader stems, root tips emerging from deeper in the soil, wider and shallower root systems, and a decrease in vertical root growth. In general, a higher PC2 score corresponds to a root system that tends to grow more narrowly near the soil surface and indicates a trade-off in the spatial arrangement of the root system, where the ability to grow deeper is constrained to individuals with a narrower root system and vice versa.

For PC3 (‘root system size’; Fig. 3C) we found that the total surface area of the root system explained most of the variation and loaded positively on this axis, therefore indicating that higher scores correspond to an overall larger root system. Multiple traits that describe overall root system morphology (*e*.*g*., root diameter and root tip count) contributed mainly to PC4 (‘root morphology’; Fig. 3D). Overall, higher scores on the morphology axis correspond to a root system that has multiple lateral roots and smaller lateral root diameter (*i*.*e*., thinner lateral roots) and one that exhibits a greater range in the rooting angles relative to the soil surface and to the tap root (*i*.*e*., develops roots that grow both obtuse and acute relative to soil surface and tap root). As such, *I. purpurea* individuals that produce many lateral roots tend to produce smaller roots with less diverse rooting angles.

### Evidence of belowground competition

We found a significant effect of treatment (*F*_1,370.05_ = 3.98, *p-value* = 0.046) with *I. purpurea* producing 18% fewer seeds when in the presence of competition with *I. hederacea* compared to growing alone. We also uncovered a significant treatment by block interaction (*F*_3,371.54_ = 2.62, *p-value* = 0.05). These results indicate that *I. purpurea* competed with *I. hederacea* belowground and that the intensity of competition was environmentally dependent.

### Maternal line variation in root traits

From our linear mixed model ANOVA on each of the PCs, we uncovered evidence for maternal line variation in root morphology (χ^2^ = 6.31, *p-value* = 0.01; Table 1) but no evidence for maternal line variation in root topology, architecture or size (Table 1). In addition, all four modular traits differed with environment (block), but not with competition (Table 1).

**Table 1.** Linear mixed model results for the modular root system traits obtained from the first four principal components within *I. purpurea. F-statistics* and χ^2^ values show the effects of Block, Treatment, and Maternal Line variation, respectively. *p-values* of fixed and random effects are reported within parentheses. Bolded values indicate *p-value* < 0.05.

| Trait | *F-statistics* | | $\chi^2$ |
| --- | --- | --- | --- |
|  | Block<br>DF = 3 | Treatment<br>DF = 1 | Maternal Line<br>DF = 1 |
| Root topology (PC1) | 1660.90<br>(2e-16) | 0.77<br>(.38) | -4.55e-13<br>(.99) |
| Root architecture (PC2) | 23.66<br>(1.51e-13) | 1.42<br>(.23) | 2.17<br>(.14) |
| Root size (PC3) | 5.04<br>(.002) | 2.32<br>(.13) | 0.48<br>(.49) |
| Root morphology (PC4) | 5.50<br>(0.001) | 0.08<br>(.78) | **6.31**<br>**(.01)** |

In addition, we performed *post hoc* linear mixed models on eight individual root morphology traits and three individual root architecture traits because we found evidence that belowground competition altered selection on these traits (see results below). This analysis uncovered significant maternal line variation for soil tissue angle range (χ^2^ = 4.66, *p-value* = 0.03; Table S3), root tissue angle range (χ^2^ = 4.22, *p-value* = 0.04; Table S3) and maximum soil tissue angle (χ^2^ = 5.17, *p-value* = 0.02; Table S3), indicating that these individual root traits can potentially respond to selection. The block effect explained a significant proportion of variation in all these specific traits while competition did not.

### Testing for selection on root traits

Selection analysis on the modular root traits showed evidence for negative directional selection on root morphology (PC4) (β = −0.17, *p-value* = 0.03; Table S3; Fig. 4) when *I. purpurea* was grown in the presence of competition, and no evidence for selection on root morphology in the absence of competition (Table S3; Fig. 4). ANCOVA revealed a significant treatment × trait interaction (*F*_*1,16*_ = 5.33, *p-value* = 0.03; Table S3), showing that the pattern of selection on root morphology was altered by belowground competition. These results indicate that belowground competition imposes selection for root systems that exhibit smaller root morphology (*i*.*e*., a decrease in overall lateral root production with an increase in lateral root diameter along with selection for a decreased range of root angles). We did not find evidence for selection on root topology, or architecture (*i*.*e*., PC1 & PC2) in either the presence or absence of competition (Table S3), suggesting that these traits are not under selection regardless of the competitive environment, or alternatively, that the signal of selection on specific traits contributing to each PC was diluted.

**Fig. 4.**
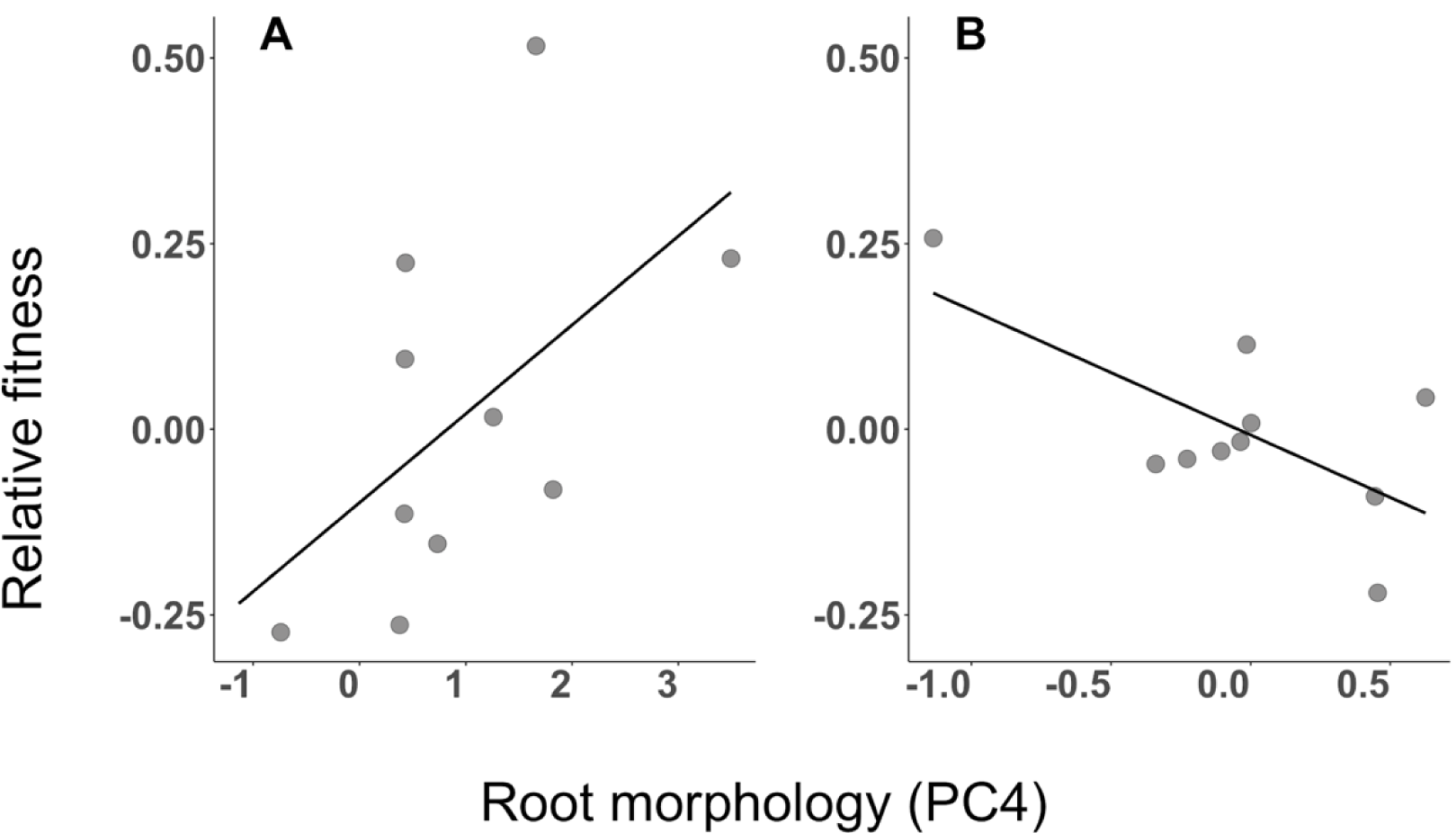
Linear regression of root morphology onto relative fitness by treatment. Root morphology was mean standardized and averaged by maternal line and treatment. There is nonsignificant positive selection on root morphology when *I. purpurea* is in the absence of competition (A) (β = 0.12, *p-value* = 0.10), and a significant negative selection (B) (β = −0.17, *p-value* = 0.03) when *I. purpurea* is in competition with *I. hederacea*. ANCOVA showed that the Treatment × root morphology is significant (*F*_1,16_ = 5.33, *p-value* = 0.03; Table S4), indicating that competition influences the pattern selection on root morphology as a modular trait.

We next evaluated evidence for selection on individual root traits via PC back regression since the absence of selection at the modular level does not necessarily reflect absence of selection on specific root traits. In the absence of competition, we uncovered positive selection on skeleton node number, root tip count, the number of adventitious roots, and negative selection on average root density (Table 2). In the presence of competition, however, we found evidence for positive selection on mean root tip diameter, hypocotyl diameter, tap root diameter and maximum diameter 90-100% soil depth, and negative selection on soil tissue angle range, maximum soil tissue angle and root tissue angle range within *I. purpurea* (Table 2). Although maximum soil root tissue angle and soil and root tissue angle range describe spatial characteristics of the root system, they contribute mainly to PC4 (root morphology) (Fig. 3; Table S1), therefore contributing to selection acting on root morphology in the presence of competition. The 95% confidence intervals on selection gradients each of these traits did not overlap between treatments, indicating that, with the exception of skeleton node number, belowground competition altered the pattern of selection on these traits.

### Test of character displacement

We found a negative linear relationship between phenotypic distance in root architecture (PC2) and relative fitness (β = −0.06, *p-value* = 0.03; Table 3; Fig. 5), suggesting that competitor individuals with similar root architectures exhibited higher fitness than competitor individuals with more divergent architectural traits (*i*.*e*., character convergence rather than displacement). We found no evidence of a linear relationship between standardized relative fitness and phenotypic distances in root topology (PC1) or morphology (PC4) (Table 3).

**Table 3.** Test for the hallmark prediction of character displacement in modular root system traits of *I. purpurea*. Phenotypic distance was calculated as the absolute Euclidean distance between competitor pairs of *I. purpurea* and *I. hederacea* for root topology (PC1), root architecture (PC2) and root morphology (PC4). Phenotypic distances were averaged by maternal Line and maternal line *×* species combination, and then regressed onto standardized values of relative fitness; ± 1 standard error is reported next to its corresponding linear regression slope (β coefficient). *p-values* are indicted within parentheses. Bolded values indicate a *p-value* < 0.05.

| Phenotypic distance<br>($\sqrt{PC_{I.purpurea}^2 - PC_{I.hederacea}^2}$) | $\beta$ coefficient |
| --- | --- |
| Root topology (PC1) | 0.03 ± 0.03 (.24) |
| Root architecture (PC2) | **-0.06 ± 0.03 (.03)** |
| Root morphology (PC4) | 0.04 ± 0.07 (.62) |

**Fig. 5.**
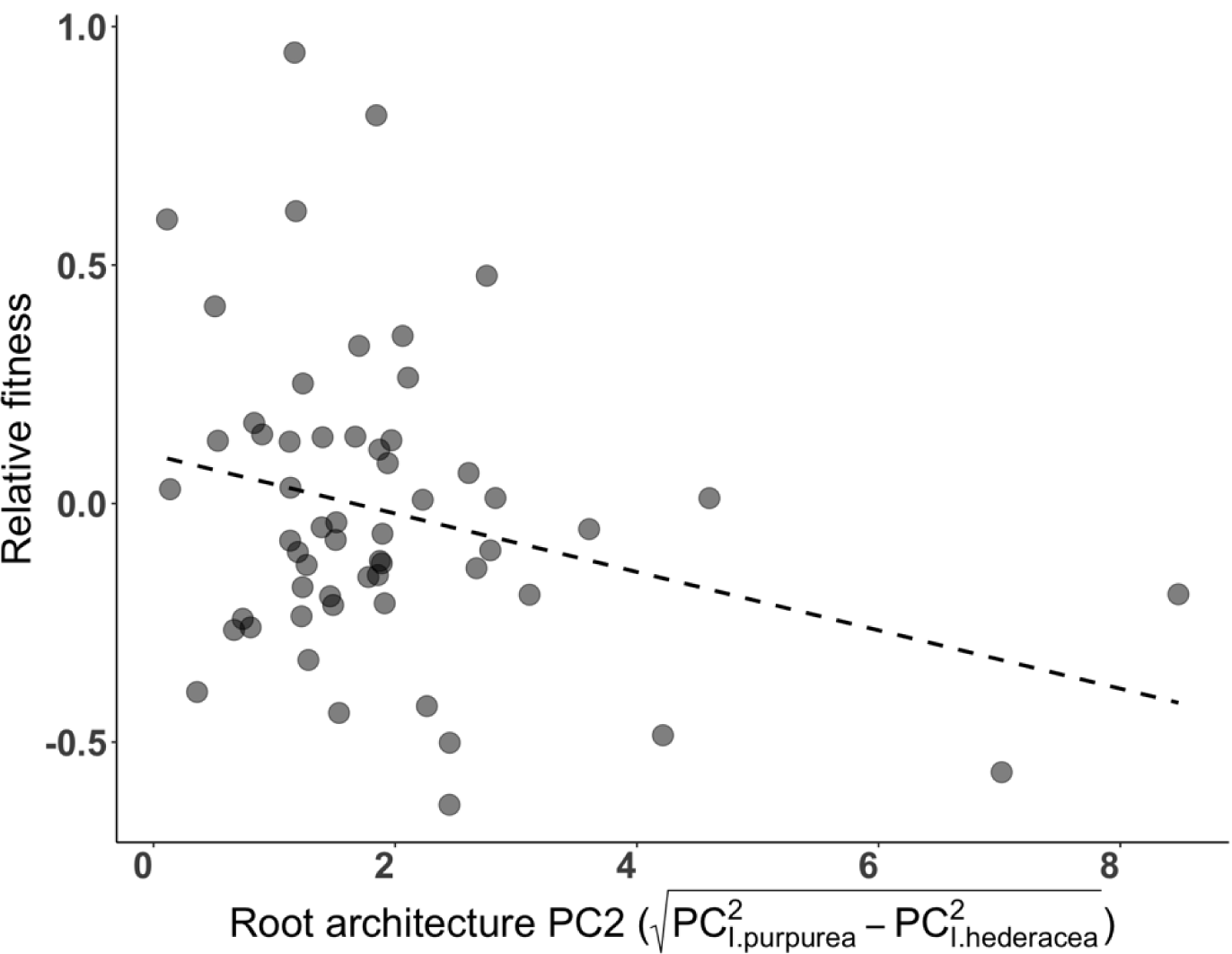
Negative relationship (β = −0.06 ± 0.03, *p-value* = 0.04; Table 4) between phenotypic distance of root architecture (PC2) and standardized relative fitness for *I. purpurea* when in competition with *I. hederacea*. The phenotypic distance of root architecture was calculated as the Euclidean distance in PC2 between competing pairs of *I. purpurea* and *I. hederacea* after the removal of Block effects, and then averaged by maternal line and species by maternal line combination type. Each point represents two to eight biological replicates.

Finally, since we found evidence that belowground competition altered selection on a handful of individual root traits (*see* Testing for selection on root traits *above*), we performed *post hoc* tests to examine the pattern of character displacement on eight individual root morphology traits and three individual root architecture traits. However, we found no evidence for a significant linear relationship between phenotypic distance between these traits and relative fitness (results not shown).

## Discussion

Our research examined the potential that root traits may evolve *via* the process of character displacement. We performed a field experiment where we grew *I. purpurea* (focal species) in the presence and absence of *I. hederacea* and determined if the process of character displacement could influence the evolution of root traits by testing four key criteria. We found that *I. purpurea* grown in the presence of *I. hederacea* experienced a significant reduction in fitness, thus providing evidence that these species compete belowground, a result that is in alignment with our previous field study (Criterion 1; Colom & Baucom, 2020). We uncovered evidence for genetic variation in the modular trait root morphology and for three individual traits, demonstrating that multiple root traits represent viable targets of selection (Criterion 2). Further, we found that belowground competition imposed selection on root morphology as a modular trait and on multiple individual root traits, indicating that belowground competition can act as an agent of selection (Criterion 3). Importantly, our test for the hallmark prediction of character displacement (Criterion 4) revealed a negative linear association between plant fitness and phenotypic distance for root architecture as a modular trait. This result did not show evidence for the potential for character displacement as we hypothesized, but instead evidence for character *convergence* in root architecture. Below, we expand on the implications of our findings and our interpretations in light of current experimental and theoretical work in root trait biology and ecology.

### Genetic variation in root traits suggests evolutionary potential

Our finding of significant maternal line variation in root morphology as a modular trait and in individual root traits shows that these traits exhibit the potential to respond to selection and evolve. These results are in line with previously reported evidence for maternal line variation in root traits associated with root system architecture and morphology in both *I. purpurea* and *I. hederacea* (Colom & Baucom, 2020). Interestingly, these previous results were based on measurements taken from individuals that were grown in greenhouse conditions, where environmental conditions are simple. That we also uncovered maternal line variation for specific root architecture traits, and root morphology as a modular trait under field conditions strengthens support for these traits as viable targets of selection that can potentially evolve given selection from belowground competition.

### Belowground competition generates selection on root traits

Although we found evidence that multiple root traits have the potential to respond to selection, evidence that interspecific competition alters the pattern of selection on these same root phenotypes is necessary for making a strong case that belowground competition can lead to character displacement. As such, we examined selection on root traits using two approaches: selection at the modular level (*i*.*e*., PC’s), and selection on specific root traits. Selection analysis on each PC is appropriate for studying the root system given that many root traits are strongly correlated (Chong *et al*., 2018). However, if traits that load strongly on a single PC axis do not contribute to fitness, the signal for selection on traits that may contribute to that PC could go undetected (MitchellLJOlds & Shaw 1987; Chong *et al*., 2018). Therefore, we also performed PC back regression, a linear algebra transformation technique that allows us to project selection gradients back into original trait space, and compute estimates of selection coefficients on specific traits (Chong *et al*., 2018). We found that directional selection acted on root morphology as a modular trait, and that the direction of selection was altered according to competitive context. Specifically, our results show a pattern of negative selection on root morphology in the presence of competition, and positive (albeit nonsignificant) selection on root morphology in the absence of competition. This result indicates that competition was selecting on thicker but fewer number of lateral roots, whereas in the absence of competition we did not uncover any evidence of selection.

One potential reason we uncovered a pattern of selection for smaller values of root morphology in the presence of competition may be due to specific foraging strategies that provide a benefit in this environment. For example, the production of fewer and thicker lateral roots has been linked with resource conservation, suggesting that selection is favoring individuals of *I. purpurea* that may acquire nutrients efficiently (Eissenstat & Yanai, 2002; Paula & Pausas, 2011). This explanation is in line with theoretical models of belowground plant-plant competition which predict that when soil resources are low, efficient root foraging phenotypes are favored over exploitative ones because the ‘per-root’ costs are high relative to resource uptake (Hutchings and John 2007; O’Brien *et al*., 2007; McNickle & Brown, 2012). Our results also suggest that belowground competition is selecting for a wider root system (*i*.*e*., decrease in the angle formed between the soil surface and a given lateral root; *see* sketch of ‘soil tissue angle’ in Fig. S1) and decrease in the rooting angle range relative to the soil surface and tap root. The finding of negative selection on these root angle traits may reflect increased competition for topsoil resources when *I. purpurea* and *I. hederacea* grow close to each other.

In previous work we found evidence for selection for shallower root systems in *I. hederacea* as a response to belowground competition from *I. purpurea*, but no evidence of selection when considering *I. purpurea* as the focal species in competition with *I. hederacea* (Baucom & Colom, 2020). The results presented here appear to contradict our previous work, however, in our present study we measured more and different architectural traits than in Colom and Baucom (2020). In our present study we found evidence of negative directional selection acting on a similar trait, the maximum angle formed between lateral roots and the soil surface when *I. purpurea* was grown in the presence of competition but not in the absence of competition. This result implies that belowground competition from *I. hederacea* is generating selection for a decrease in the maximum rooting angle formed across lateral roots relative to the soil surface, or a shallower root system. Collectively, our past and present results indicate that root architecture plays an important role in how these plants compete and access to belowground resources and it indicates that competition for topsoil resources is strong between these two *Ipomoea* species.

Consistent with our results for selection on root morphology at the modular level, PC back regression revealed evidence that belowground competition altered selection on traits that contribute mainly to this axis, including: soil root tissue angle range, maximum soil root tissue angle and root tissue angle range and multiple root diameter and lateral root number traits (Table S2). Therefore, selection on these individual root traits are driving the patterns of selection observed at the modular level. In contrast, we did not detect evidence for selection on specific root traits that contributed to root topology or architecture at the modular level, which is consistent with the lack of evidence for selection on modular root topology and architecture.

### Character convergence but not displacement on root traits

Traditional hypotheses of character displacement predict that when two co-occurring, closely related species overlap in their resource associated traits, selection should favor divergence as that would lead to lower resource overlap between species (Losos, 2000; Pfennig & Pfennig, 2009). Consequently, we predicted that fitness should increase with increasing phenotypic distance to a competitor (Criterion 4). However, we actually found the reverse result, indicating evidence for character *convergence* rather than displacement. If root architecture influences soil exploration and what resources are readilyaccessible, why did we find support for character convergence instead of divergence?

For plants, phenotypic plasticity in root architecture has been argued to represent an adaptive strategy that allows plants to access and compete for key nutrients, and further, root architecture has been shown to respond plastically to nutrient availability across multiple plant species (Fitter *et al*., 1991; Nicotra & Davidson, 2010; Yu *et al*., 2014). Therefore, one plausible explanation behind our result of character convergence in root architecture is that *I. purpurea* individuals capable of recognizing and responding plastically to both their immediate resource environment and to the presence of a competitor individual would be able to maximize fitness, whereas individuals less capable of sensing and responding to these environmental constraints would exhibit lower fitness. Soils are complex and heterogeneous, and plants may be selected to respond plastically to very local soil conditions. If both *I. purpurea* and I. *hederacea* benefit from similar plastic responses to a given local soil environment, we might expect to see patterns of trait convergence associated with higher fitness. In short, the local environment may constrain morning glories into expressing convergent phenotypes. Such constraints operating on behavior and morphology are well known from studies of competition among animal species (Gibson, 1980; Hunter & Willmer, 1989; Hunter *et al*., 1997).

Moreover, it is well established that plant root growth can respond to the presence of competitors (Cahill *et al*., 2010), with neighbor recognition hypothesized to be due to either sensing of root exudates (Bais *et al*., 2006; Biedrzyckie *et al*., 2010; *reviewed in* Pierik *et al*., 2013; Semchenko & Lepik 2014) or more simply from feedback given the internal nutrient status of the plant (*reviewed in* Pierik *et al*., 2013; McKnickle & Brown, 2014). Consistent with the idea that individuals can respond to the presence of a neighbor, supplementary analysis showed that the root architecture of *I. purpurea* varied depending on the presence of specific *I. hederacea* competitors after the removal of block effects (Table S2-5). Additionally, we found preliminary evidence for a negative linear trend between plant size and phenotypic distance in root architecture within *I. hederacea* (Fig. S2-5). Given that plant size is often a strong correlate of fecundity (Aarssen & Taylor, 1992), this result suggests that the pattern of convergence is potentially present in both species, and perhaps that both species may modify their root architecture to compete for varying limiting resources.

Phenotypic plasticity of root traits can have important implications for the evolution of the belowground root system. For example, phenotypic plasticity can obscure selection from acting on phenotypes that are genetically variable, and hence, impede traits from *responding* to selection. However, reaction norms of functional traits can be genetically variable, and therefore, phenotypic plasticity itself can represent an important target of selection that can evolve in response to different environmental stressors (*e*.*g*., competition; Via & Lande, 1985; Schlichting, 1986; Scheiner, 1993). Testing whether plasticity in root traits is a viable target of selection and whether belowground competition-imposed selection on phenotypic plasticity in root traits was beyond the scope of our current research. However, literature in the field of plant breeding has demonstrated that plasticity in root architecture can be genetically variable (*reviewed in* Jung & McCouch, 2013), indicating that phenotypic plasticity in root architecture is a viable target of selection. Whether it is possible that belowground plant-plant competition can promote the evolution of phenotypic plasticity in root architecture remains an elusive and unaddressed question.

### Conclusion

Our study shows that belowground competition results in a pattern of character convergence rather than divergence in root architecture between competing pairs of *I. purpurea* and *I. hederacea*. Moreover, we found evidence to suggest that root architecture responds plastically to its specific competitive environment, which may reflect an adaptive mechanism that allows plants to compete for multiple key nutrients. Additional research will be required to assess whether phenotypic plasticity in root architecture can potentially evolve as a response to belowground competition and result in patterns of character displacement or convergence in plasticity. Our work emphasizes the importance of belowground competition to potentially influence the evolution of the root system and considers its complex and integrated nature. Therefore, we encourage other researchers to examine the potential for character convergence/displacement in different study systems/environments, and further, to consider phenotypic plasticity as a target of selection. Future work and experimental replication will allow us to understand how widespread and viable these evolutionary processes are in nature.

## Data availability statement

The R code is available at https://github.com/SaraMColom/CharacterDisplacement, and the data will be uploaded to the Dryad Digital Repository.

## Acknowledgments

We graciously thank Deborah Goldberg, Mark Hunter and members of the Baucom Lab for their helpful comments on earlier versions of this article. We thank John Stinchcombe for his advice on statistical analysis, and Alexander Bucksch for his assistance in interpreting the root traits quantified in our study. We thank Andres Ibarra, Jazlyn Marcos, Yoav Jacob and Donàa Williams for their invaluable assistance with planting, maintenance of the field site, and sample collection and processing. We thank employees of Matthaei Botanical Gardens, especially Michael Palmer, Paul Girard and Jeremy Moghtader for their expertise and loan of field equipment and machinery. This work was made possible with financial support of internal grants at the University of Michigan.

## Supplementary Figures & Tables

**Table S1.** lists the full root trait name and its corresponding abbreviated name (‘Code’) assigned by DIRT software for the 33 traits that were analyzed in the study. Each root trait was cataloged into four functional trait classes indicated within parenthesis *a priori*. We report the PC axis where a given individual root trait contributed the most variance and report its percent contribution to that axis (‘PC Contribution’). Traits with ‘*’ indicate multiple traits and a corresponding range in their percent (%) contribution to a given PC axis.

| **Trait name** | **Functional Class** | **Description** | **Trait Code** | **PC Contribution (%)** |
| --- | --- | --- | --- | --- |
| Accumulated width* | Topology | Percentage of width accumulation at 10-90 % depth | D10-D90 | PC1: 7.00-10.69 % |
| Skeleton width | Architecture | Width calculated from the medial axis of the root system. | SKL WIDTH | PC2: 7.50 % |
| Skeleton depth | Architecture | Depth calculated from the medial axis of the root system. | SKL DEPTH | PC2: 10.84 % |
| Maximum soil tissue angle | Architecture | Maximum soil tissue angle measured over all root tips. | STA MAX | PC4: 6.94 % |
| Maximum width of the root system | Architecture | Maximum root system width measured from first to last horizontal foreground pixel. | WIDTH MAX | PC2: 7.53 % |
| Maximum root tissue angle | Architecture | Maximum root tissue angle measured over all root tip paths. | RTA MAX | PC2: 9.02 % |
| Root top angle range | Architecture | Range of root tissue angles present in the root system. | RTA RANGE | PC4: 5.73 % |
| Spatial root distribution X | Architecture | Spatial distribution of the root shape in the x-axis. This is the x component of the vector pointing from the center of the bounding box of the root shape to the center of mass of the root shape. | RDISTR X | PC2: 0.22 % |
| Spatial root distribution Y | Architecture | Spatial distribution of the root shape in the y-axis. This is the y component of the vector pointing from the center of the bounding box of the root shape to the center of mass of the root shape. | RDISTR Y | PC2: 8.01 % |
| Adventitious root angle | Architecture | Adventitious root angle estimated from the paths detected in the number of adventitious roots. | ADVT ANG | PC2: 4.83 % |
| Basal root angle | Architecture | Basal root angles estimated from the paths detected in the number of basal roots. | BASAL ANG | PC2: 6.61 % |
| 50 percent drop | Architecture | Depth value where 50% of the root tip paths emerged from the central path. | DROP 50 | PC2: 10.85 % |
| Projected root area | Size | Number of foreground pixels belonging to the root system. | AREA | PC3: 13.49 % |
| Skeleton nodes | Morphology | Nodes calculated from the medial axis of the root system. | SKL_NODES | PC4: 8.25 % |
| Stem diameter | Morphology | Stem diameter derived from medial axis. | DIA STM | PC3: 2.31 % |
| Simple stem diameter | Morphology | Simple stem diameter calculated in root estimator for shovelomics | DIA STM SIMPLE | PC2: 11.51 % |
| Average root density | Morphology | Ratio of foreground to background pixels within | AVG DENSITY | PC4: 1.43 % |
| Mean tip diameter | Morphology | Mean tip diameter estimated from the medial circle at the tips. | TD AVG | PC4: 14. 53 % |
| Root tip count | Morphology | Overall number of tips detected in the image | RTP COUNT | PC3: 9.37 % |
| Number of Adventitious | Morphology | Number of root tip paths emerging from root segment 1. | ADVT COUNT | PC4: 3.47 % |
| Number of basal roots | Morphology | Number of basal roots estimated as emerging root tip bundles from root segment 2. | BASAL COUNT | PC4: 5.64 % |
| Basal root angle | Architecture | Basal root angles estimated from the paths detected in the number of basal roots. | BASAL ANG | PC2: 6.61 % |
| Hypocotyl diameter | Morphology | Hypocotyl diameter estimated over detected hypocotyl region as the average diameters of medial circles. | HYP DIA | PC4: 8.46 % |
| Tap root diameter | Morphology | Tap root diameter estimated over detected taproot region as the average diameters of medial circles. | TAP DIA | PC4: 10.97 % |
| Maximum diameter at 90-100% percent depth | Morphology | Maximum diameter found in the interval of 90-100% rooting depth | MAX DIA 90 | PC4: 9.42 % |

**Fig. S1.**
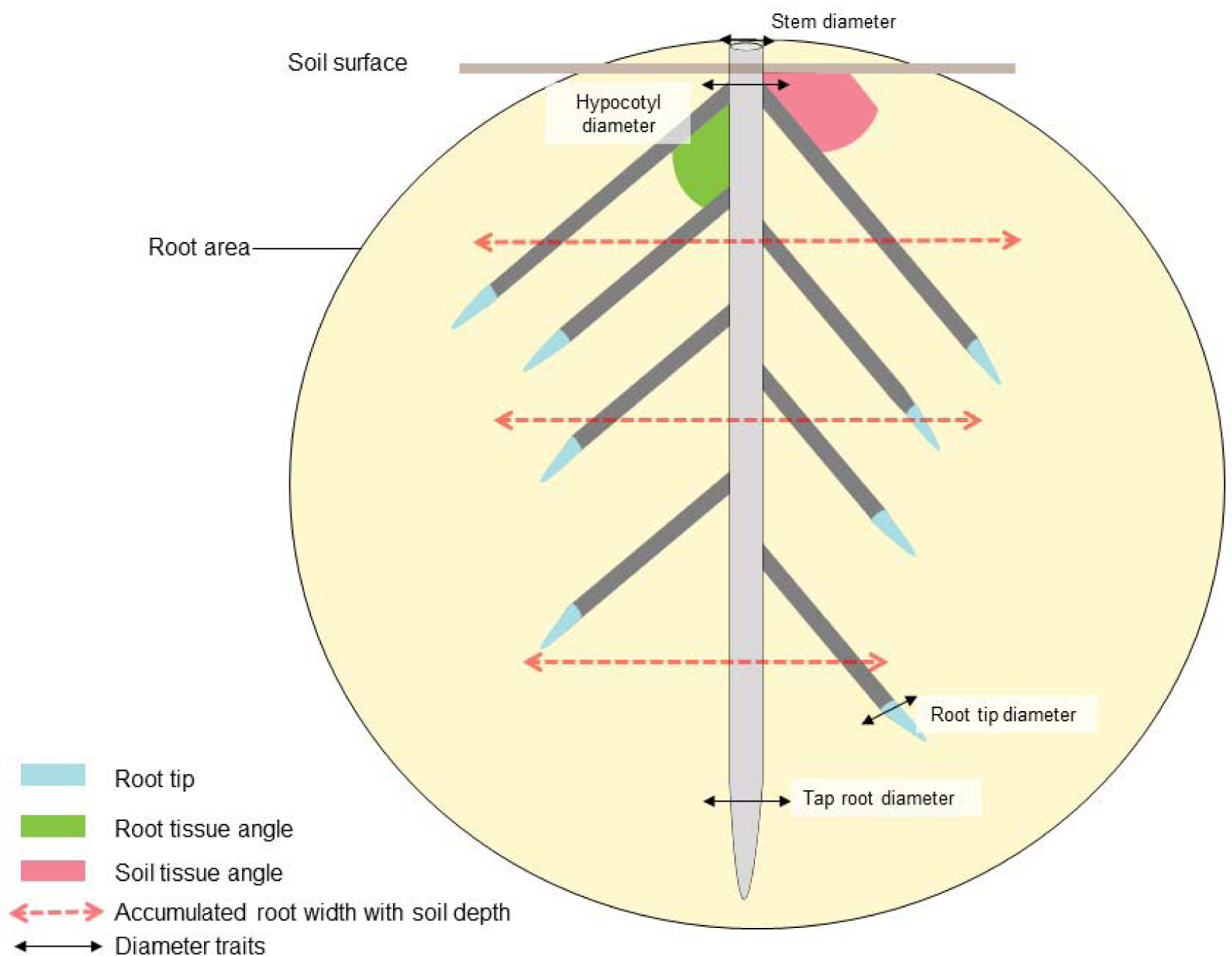
Sketch for some root traits captured by DIRT. Root area represents the ‘AREA’ trait, estimated as the total number of pixels of the root system as indicated by a light yellow circle encapsulating the entire sampled root system. Accumulated root width with soil depth is captured with the two-way dashed light red arrows (‘D%’; Table S1). Different diameter traits are indicated with two-way solid black arrows including, tap root diameter (‘TAP_DIA’), root tip diameter (‘TD_AVG’; Table S1), stem diameter (‘DIA_STEM’; Table S1) and hypocotyl diameter (‘HYP_DIA’; Table S1). Root tips are colored in light blue, and an example of soil root tissue angle and root tissue angle are shown by the pink and green shaded regions, respectively.

**Table S3.**
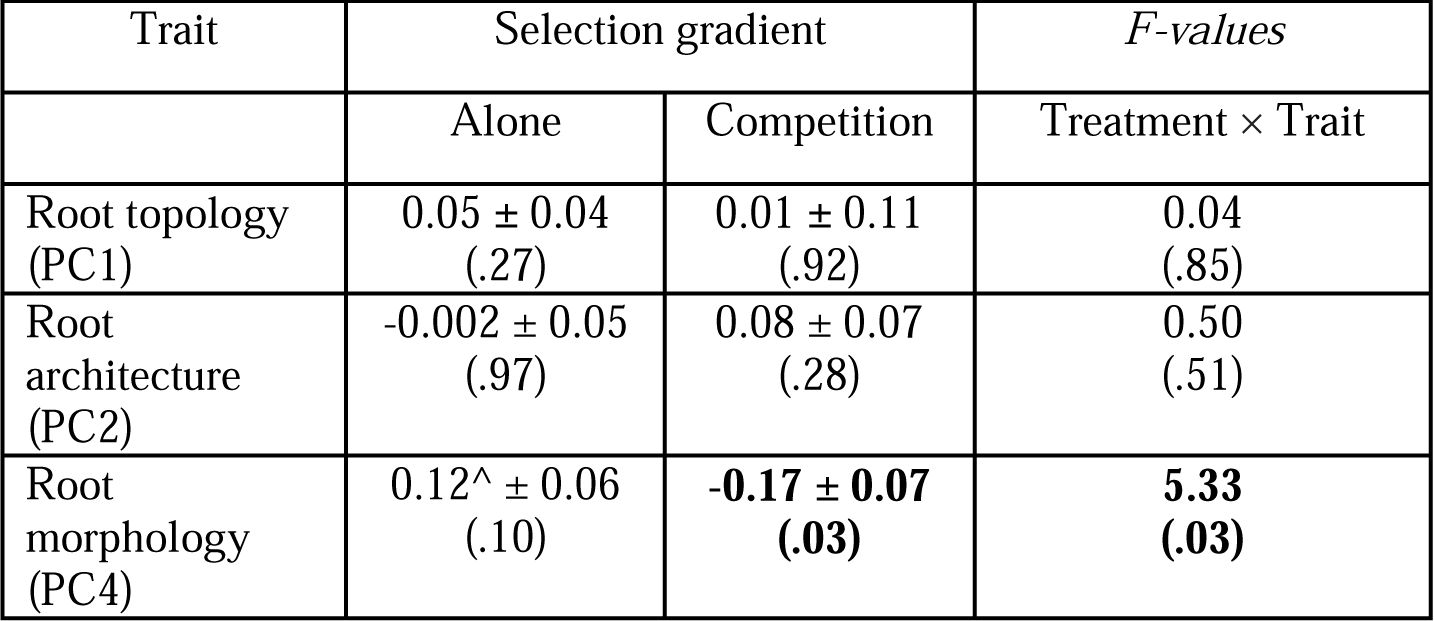
Results of linear selection gradient for root topology (PC1), architecture (PC2), and morphology (PC4), for each Treatment is presented with their respective linear regression slopes (β), its corresponding **±** 1 standard error and *p-value* in parentheses. *F-values* and their corresponding *p-values* from ANCOVA are also shown to indicate the effect of Treatment on the pattern of selection for each trait (Treatment × Trait). Bolded values indicate a *p-value* < 0.05 and ‘^’ indicates a marginally significant *p-value* = 0.10.

| Trait | Selection gradient |  | *F-values* |
| --- | --- | --- | --- |
|  | Alone | Competition | Treatment × Trait |
| Root topology (PC1) | 0.05 ± 0.04<br>(.27) | 0.01 ± 0.11<br>(.92) | 0.04<br>(.85) |
| Root architecture (PC2) | -0.002 ± 0.05<br>(.97) | 0.08 ± 0.07<br>(.28) | 0.50<br>(.51) |
| Root morphology (PC4) | 0.12<sup>^</sup> ± 0.06<br>(.10) | **-0.17 ± 0.07**<br>**(.03)** | **5.33**<br>**(.03)** |

**Fig. S2.**
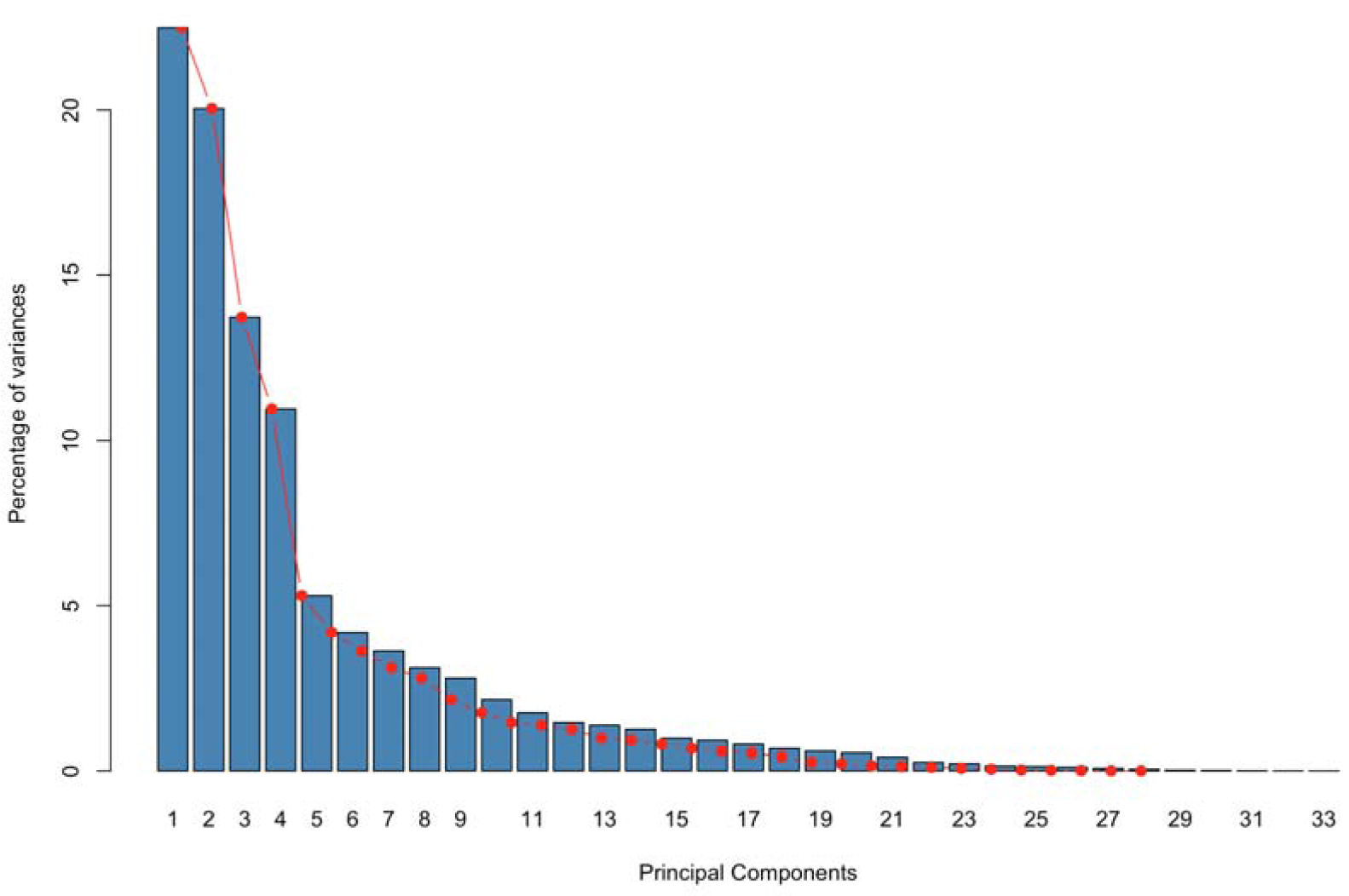
Scree plot demonstrating the percentage of variance explained by each principal component computed on the correlation matrix of the transformed root traits after the removal of Block effects. The first four principal components contribute to more than 10.0% of the total variation.

**Table S5.** Results for post hoc linear mixed model analysis on a subset of individual root traits. *F-values* and their corresponding *p-values* from ANOVA are also shown to indicate the effect of Block and Treatment on original root traits. Bolded values indicate a *p-value* < 0.05 and ^ indicates *p-value* < 0.07.

| Traits | *F*-statistics | | $\chi^2$ |
| --- | --- | --- | --- |
| Trait description | Block<br>DF = 3 | Treatment<br>DF = 1 | Maternal Line<br>DF = 1 |
| Skeleton nodes (morphology) | **130.59 (<.001)** | 2.13<br>(.15) | 0.47<br>(.50) |
| Maximum diameter at 90-100% percent depth<br>(morphology) | **1011.34 (<.001)** | 1.62 (.20) | <0.001<br>(.99) |
| Average root density (morphology) | **113.94 (<.001)** | 0.02<br>(.88) | 0.13<br>(.72) |
| Mean tip diameter (morphology) | **1208.57 (<.001)** | 2.09<br>(.15) | <0.001<br>(.99) |
| Root tip count (morphology) | **13.60 (<.001)** | 1.31<br>(.25) | 1.66<br>(.20) |
| Soil tissue angle range (architecture) | **5.70**<br>**(.001)** | 2.78<br>(.10) | **4.66**<br>**(.03)** |
| Maximum soil tissue angle (architecture) | **7.33 (<.001)** | 2.36 (.13) | **5.17 (.02)** |
| Root top angle range (architecture) | 2.29<br>(.08) | 1.82 (0.18) | **4.22**<br>**(.04)** |
| Number of Adventitious roots (morphology) | **1.29**<br>**(.28)** | 0.01 (0.91) | <0.001<br>(.99) |
| Hypocotyl diameter (morphology) | **1116.61 (<.001)** | 3.47 (.06)^ | <0.001<br>(.99) |
| Tap root diameter (morphology) | **840.17 (<.001)** | 0.52<br>(.47) | <0.001<br>(.99) |

**Table S5.** Supplementary results for linear model ANOVA on root topology, architecture and morphology to evaluate the effect of Competitor and Combination Pairing on root architecture standardized for Block effects. Root trait was treated as the response variable and Competitor and Combination Pairing were fixed effects, we ran separate models for each root trait (root topology, architecture, size and morphology), respectively. *F-values* and their corresponding *p-values* are shown to indicate the effect of Combination Pairing and Treatment on root architecture. Bolded values indicate a *p-value* < 0.05.

| Root trait | Competitor<br>DF = 5 | Combination Pairing<br>DF = 54 |
| --- | --- | --- |
| Root topology (PC1) | 1.55 (.17) | 1.33 ^ (.07) |
| Root architecture (PC2) | 2.09^ (.07) | **1.38 (.04)** |
| Root size (PC3) | **2.55 (.03)** | **1.75 (.001)** |
| Root morphology (PC4) | 0.65 (.66) | **1.47 (.02)** |

**Fig. S4.**
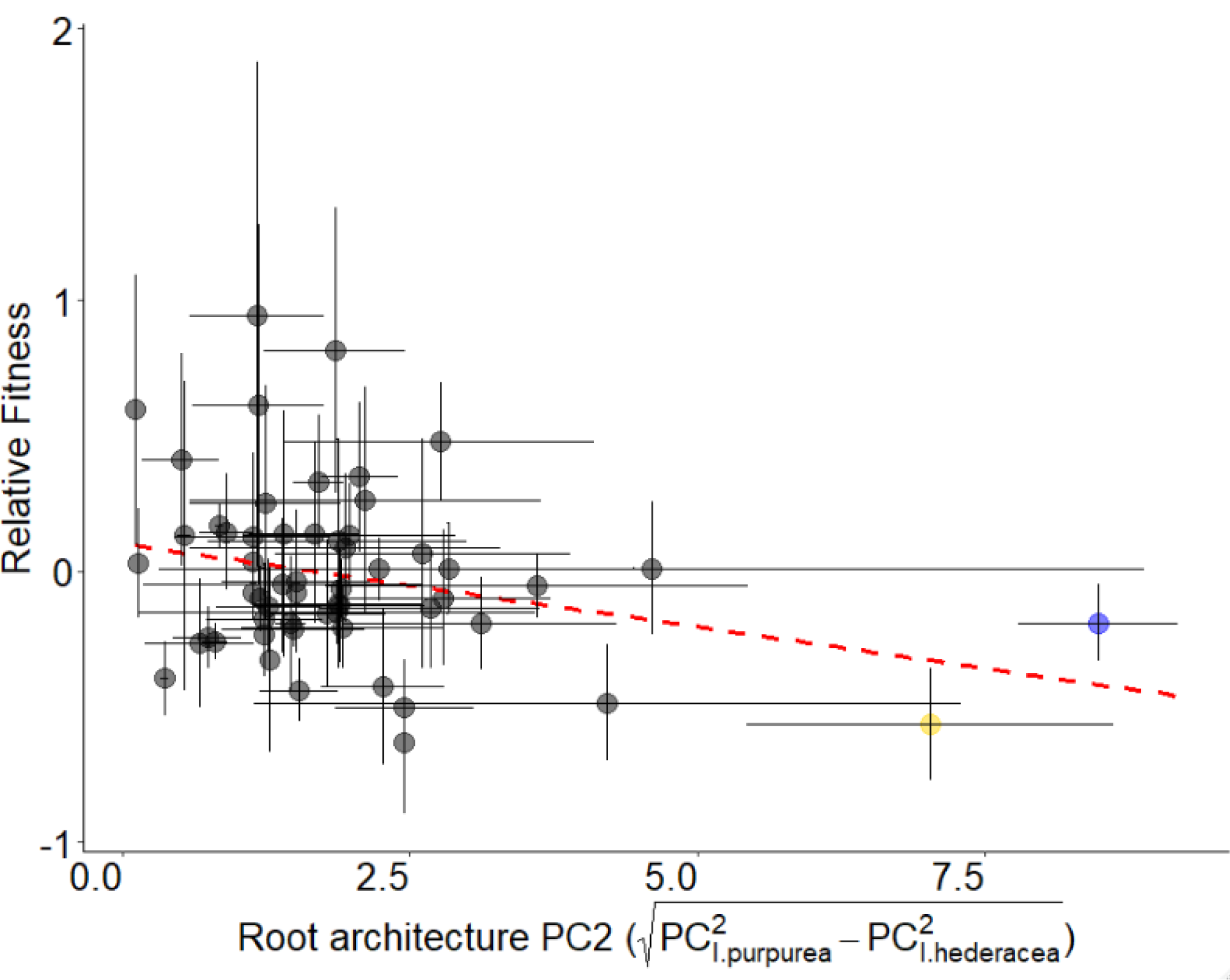
Negative relationship (β = −0.06 ± 0.03, *p-value* = 0.04; Table 4) between phenotypic distance of root architecture (PC2) and standardized relative fitness for *I. purpurea* when in competition with *I. hederacea*. The phenotypic distance of root architecture was calculated as the Euclidean distance in PC2 between competing pairs of *I. purpurea* and *I. hederacea* after the removal of Block effects, and then averaged by maternal line and species by maternal line combination type. Each point represents two to eight biological replicates. For each point, error bars were drawn based on observed values of relative fitness (Y-axis) and root architecture (X-axis) for a given maternal line within competition treatment, respectively. **Colored points (yellow and blue) indicate two outliers that were maintained in our final analysis because of their low intraspecific variation**.

**Fig. S5.**
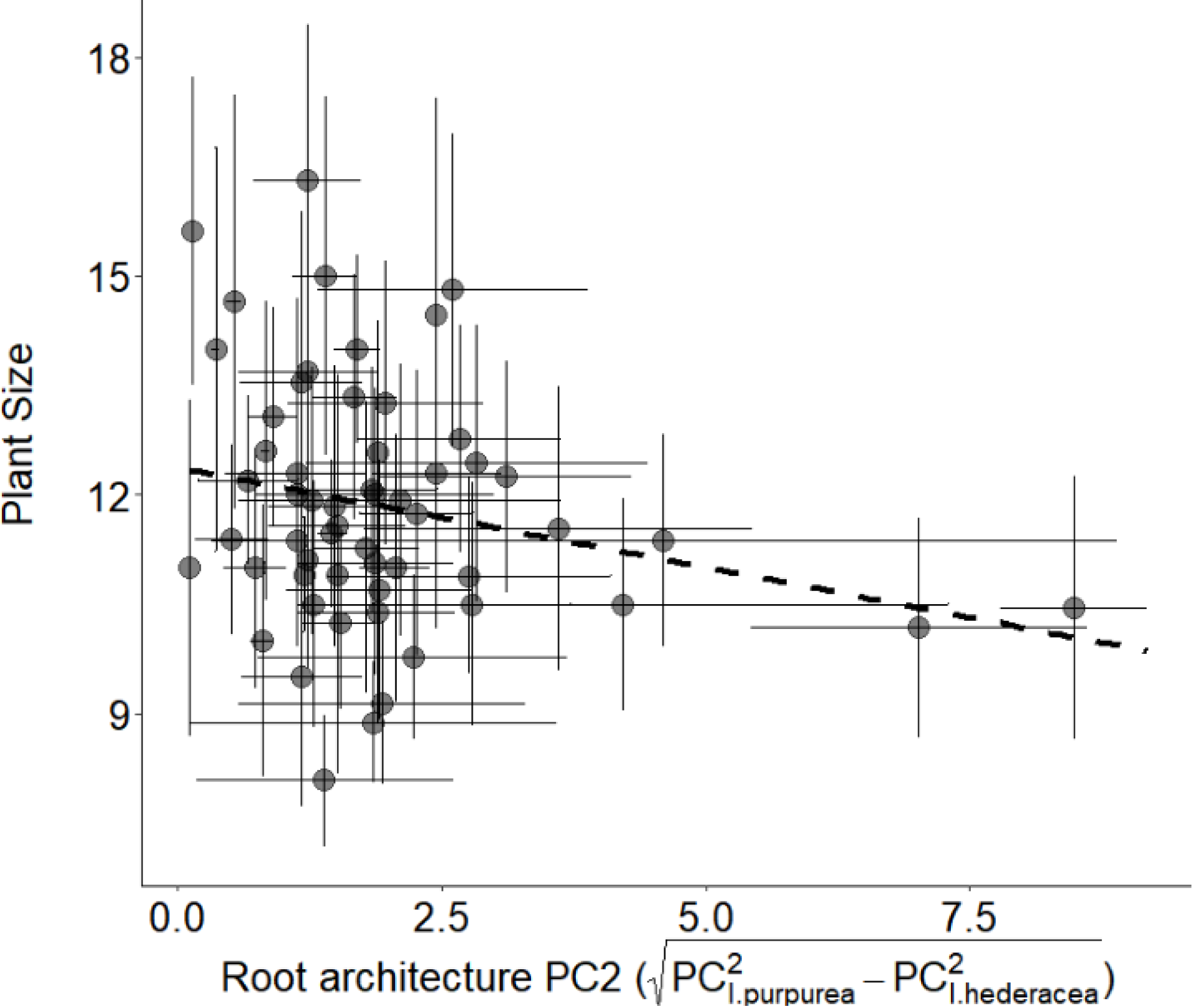
Evidence of a marginally significant negative linear relationship (β = −0.27 ± 0.15, *p-value* = 0.07) between phenotypic distance of root architecture (PC2) and plant size (*i*.*e*., leaf number averaged by combination pairing and maternal line) for *I. hederacea* when in competition with *I. purpurea*. The phenotypic distance of root architecture was calculated as the Euclidean distance in PC2 between competing pairs of *I. purpurea* and *I. hederacea* after the removal of Block effects, and then averaged by maternal line and species by maternal line combination type. Each point represents two to eight biological replicates. For each point, error bars were drawn based on observed values of relative fitness (Y-axis) and root architecture (X-axis) for a given maternal line within competition treatment, respectively, for *I. hederacea*.

